# Freshwater Snails in Schistosomiasis Endemic Communities in Ovia South-West Local Government Area in Edo State, Nigeria

**DOI:** 10.1101/2025.03.07.642031

**Authors:** E.E Adeyemi, M.S.O Aisien, E.E Enabulele

## Abstract

**Background:** Malacological surveys are crucial for determining vectors transmitting the *Schistosoma* species responsible for schistosomiasis in endemic communities. Ovia South-West Local Government Area of Edo State in Nigeria is one of the localities endemic for urogenital schistosomiasis. However, there is no report of the snail host of the disease in any community.

**Method:** A year-long freshwater snail survey was conducted in seven localities (Aden, Igbogor, Ikoha, Okopon, Okponha, Siluko, and Ugbogui) in Ovia South-West to determine the snail diversity and identify species that are intermediate hosts for schistosomiasis.

**Result:** A total of 468 snails, representing species in four genera (*Bulinus, Lanistes, Melanoides*, and *Radix*) were collected. Based on the morphological appearances of shells, five snails, *B. forskalii, B. globosus*,

*B. truncatus, L. varicus*, and *M. tuberculata* were identified at the species level. A snail, *Radix*, could not be identified at the species level. *Melanoides tuberculata* was the most abundant snail and the only species identified in all sampled locations (n = 284, 60.68%). The next abundant species was *Radix* sp. (n = 106, 22.65%), followed by *B. globosu*s (n = 42, 8.97%). The least abundant snail was *B. truncatus* (n = 5, 1.07%). At Siluko, all species of snails were present, and only *M. tuberculata* in Ugbogui. In both dry and wet seasons, all species of snails were present (wet, n = 297) (dry, n = 171). Similar snail diversity occurred in each sampling location in both seasons, with more individuals of each species collected during the wet season. The only infected snail was a single *Radix* sp. in Siluko that shed brevifurcate-apharyngeate distome furcocercariae. All other snail species were uninfected.

**Conclusion:** For the first time in Edo State, snail vectors of schistosomiasis were reported but were uninfected with *Schistosoma* parasite. The brevifurcate-apharyngeate distome furcocercariae from the *Radix* sp. had conspicuous eye spots that suggest it is not a *Schistosoma* sp.

## Introduction

Aquatic snails are part of the invertebrate fauna that provide ecosystem services (Strong *et al*., 2007). Besides their benefits, some freshwater snails are intermediate hosts for several parasites of public health and veterinary importance. Their presence, distribution, and abundance have implications for the epidemiology and outbreak of parasitic diseases (Pathak *et al*., 2023; Gaye *et al*., 2024). Two genera of snails, *Biomphalaria* and *Bulinus*, are of public health importance in Africa because of their roles as vectors of schistosomiasis (Brown, 1994).

Besides *Biomphalaria* and *Bulinu*s species, other reported freshwater snails in Nigeria include *Amerianna carinatus, Gabbiella africana, Gyraulus costulatus, Lanistes libycus, L. ovum, L. varicus, Lymnaea natalensis* (*Radix natalensis*), *Melanoides tubercula*ta, *Pila wernei, P. ovata, Physella acuta, Ph. marmorat*a, *Ph. waterloti, Potadoma freethi, Po. liberiensis, Po. moerchi, Indoplanorbis exustus*, and *Segmentorbis augustu*s (Ndifon and Ukoli, 1989; Ofoezie, 1999; Oloyede *et al*., 2016; Oladejo *et al*., 2021; Oso and Odaibo, 2021; Qadeer *et al*., 2023).

In Edo State, southern Nigeria, most studies have prioritized the prevalence of schistosomiasis without information on the malacological aspect of the disease transmission (Adeyemi *et al*., 2014; Noriode *et al*., 2017; Enabulele *et al*., 2021). Before this study, only Igbinosa *et al*. (2015), in a six-month survey in four localities in Ovia South-West Local Government Area of Edo State, reported three freshwater snails, *Gabbiella humerosa, L. varicus*, and *M. tuberculata*. This present study aimed to conduct a year-long malacological survey in seven schistosomiasis endemic communities (Aden, Igbogor, Ikoha, Okopon, Okponha, Siluko, and Ugbogui) in Ovia South-West LGA of Edo State to determine the snail diversity and identify species that are intermediate hosts for schistosomiasis.

## Materials and Methods

### Study area

Ovia South–West is one of the 18 Local Government Areas (LGA) of Edo State in Nigeria (Figure 1A). The LGA occupies an area between coordinates 6.4653° N and 5.3103° E. Two seasons, dry (October to March) and wet (April - September) are regular patterns in the locality. We conducted malacological surveys in seven communities with high urogenital schistosomiasis prevalence identified in our previous publication (Adeyemi *et al*., 2014). The communities sampled for freshwater snails were Aden, Igbogor, Ikoha, Okopon, Okponha, Siluko, and Ugbogui (Figure 1B).

**Figure 1.**
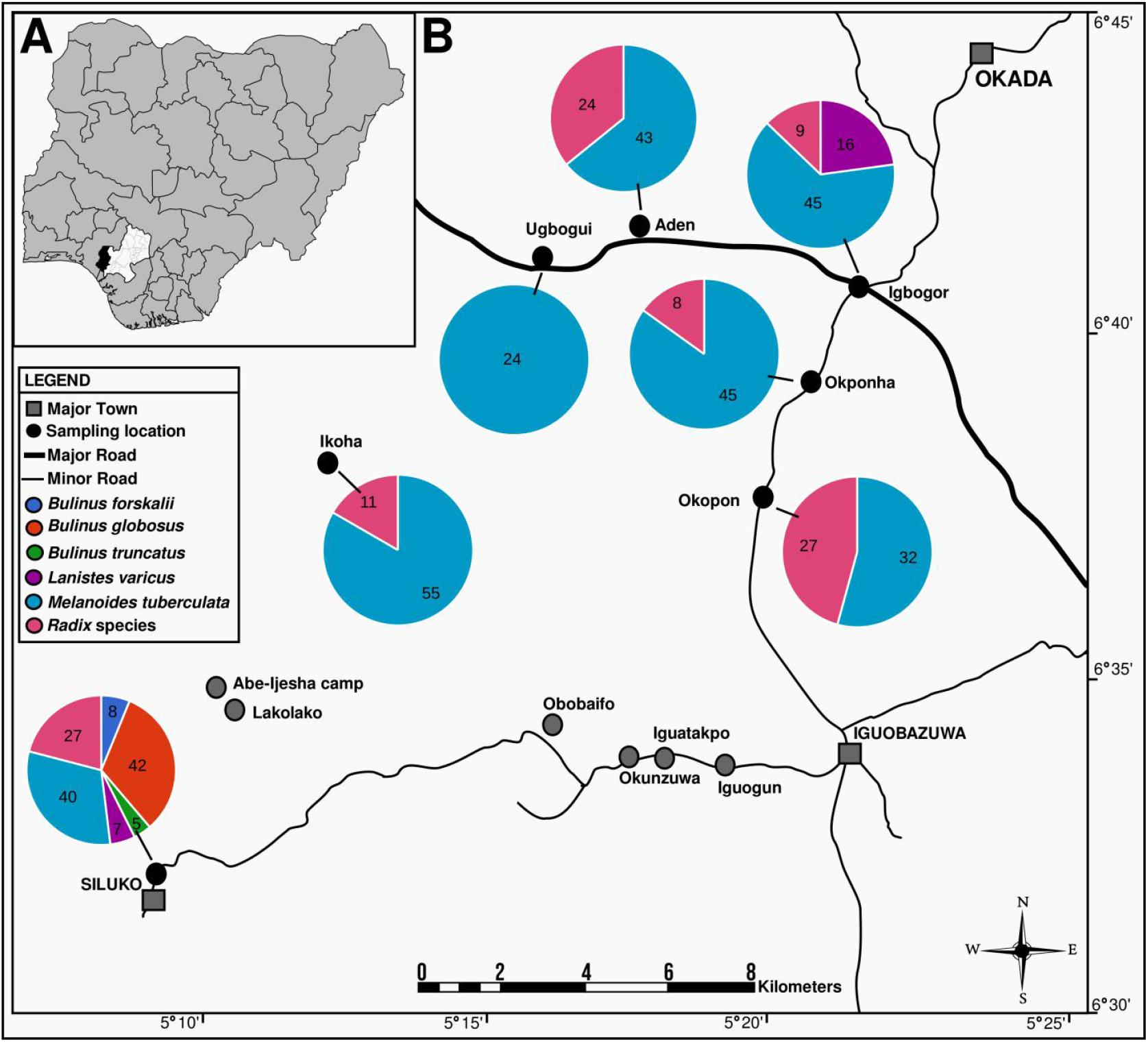
Map of snail sampling locations in Ovia South-West Local Government of Edo State. A, insert map of Nigeria. Edo state is highlighted in white and Ovia South-West is highlighted in black. B, snail sampling locations. Pie chart shows the number of snail species recorded in each location.

### Snail sampling and shedding for trematode infection

Sampling of snails was conducted in permanent water bodies (streams and rivers) between the hours of 10.00 and 14.00 from June 2009 to October 2010. Sampling was random for 30 minutes in sites where human-water contacts were highest. At each sampling location, snail collection was done with a scoop sieve and handpicking for individuals attached to vegetation or debris. The snails were transported to the laboratory for sorting, identified, and counted for abundance and relative abundance. Snail shell morphology was used for identification (Brown, 1994). Alpha diversity metrics, including Dominance, Shannon, Simpson, and Evenness, were calculated using PAST v.5.0.2 software (Hammer *et al*., 2001). The statistical significance of snail diversity across the sampling locations and between dry and wet seasons was determined using non-parametric tests Kruskal-Wallis and Mann-Whitney U.

Before shedding snails for trematode infection, they were separated by species into containers, then placed in a dark cupboard overnight and shed the following day. Shedding for each snail was in a Petri dish containing tap water, exposed to light for six hours, and visually checked for cercariae. Emerging cercariae were individually isolated on microscope slides, photographed, and morphologically identified according to the identification keys by Frandsen and Christensen (1984).

## Results

### Snail Distribution and Abundance

Four hundred and sixty-eight snails representing species in four genera (*Bulinu*s, *Lanistes, Melanoides*, and *Radix*) were collected from rivers in seven different localities in Ovia South-West LGA (Table 1). Based on the morphological appearances of shells, five snails, *B. forskalii, B. globosus, B. truncatus, L. varicus*, and *M. tuberculat*a were identified at the species level. One snail could only be identified to the genus level of classification as *Radix* sp. (Figure 2B). *Melanoides tuberculat*a was the most abundant snail and the only species recorded in all sampled locations (n = 284, 60.68%). The next abundant species was *Radix* sp. (n = 106, 22.65%), followed by *B. globosus* (n = 42, 8.97%). The least abundant snail was *B. truncatus* (n = 5, 1.07%). All six snail taxa were recorded only in Siluko, and three species (*L. varicus, M. tuberculata*, and *Radix* sp.) in Igbogor, Aden, Ikoha, Okopon, and Okponha recorded two snail species (*M. tuberculata* and *Radix* sp.). *Melanoides tuberculata* was the lone species collected in Ugbogui (Table 1). In both seasons, all taxa of snails were present, and more individuals were collected in the wet season (n = 297) than in the dry season (n = 171) (Table 2). In each sampling location, snail diversity was the same in both seasons, with more individuals collected during the wet season (Figure 2A).

**Table 1.** Summary of the distribution and abundance of freshwater snails collected in Ovia South-West, Edo State, Nigeria.

| Species | Sampling locations |  |  |  |  |  |  | Total of species | % Abundance |
| --- | --- | --- | --- | --- | --- | --- | --- | --- | --- |
|  | Aden | Igbogor | Ikoha | Okopon | Okponha | Siluko | Ugbogui |  |  |
| *Bulinus forskalii* | 0 | 0 | 0 | 0 | 0 | 8 | 0 | 8 | 1.71 |
| *Bulinus globosus* | 0 | 0 | 0 | 0 | 0 | 42 | 0 | 42 | 8.97 |
| *Bulinus truncatus* | 0 | 0 | 0 | 0 | 0 | 5 | 0 | 5 | 1.07 |
| *Lanistes varicus* | 0 | 16 | 0 | 0 | 0 | 7 | 0 | 23 | 4.91 |
| *Melanoides tuberculata* | 43 | 45 | 55 | 32 | 45 | 40 | 24 | 284 | 60.68 |
| *Radix* sp. | 24 | 9 | 11 | 27 | 8 | 27 | 0 | 106 | 22.65 |
| Number of species | 2 | 3 | 2 | 2 | 2 | 6 | 1 |  |  |
| Total snails | 67 | 70 | 66 | 59 | 53 | 129 | 24 | 468 |  |

**Table 2.** Diversity indices of freshwater snails collected in Ovia South-West, Edo State, Nigeria.

| Indices | Sampling locations |  |  |  |  |  |  | Dry Season | Wet Season |
| --- | --- | --- | --- | --- | --- | --- | --- | --- | --- |
|  | Aden | Igbogor | Ikoha | Okopon | Okponha | Siluko | Ugbogui |  |  |
| Taxa (S) | 2 | 3 | 2 | 2 | 2 | 6 | 1 | 6 | 6 |
| Individuals | 67 | 70 | 66 | 59 | 53 | 129 | 24 | 171 | 297 |
| Dominance (D) | 0.533 | 0.475 | 0.718 | 0.495 | 0.739 | 0.248 | 1 | 0.464 | 0.409 |
| Simpson (1-D) | 0.467 | 0.526 | 0.282 | 0.505 | 0.261 | 0.752 | 0 | 0.536 | 0.591 |
| Shannon (H) | 0.660 | 0.899 | 0.458 | 0.698 | 0.434 | 1.532 | 0 | 1.073 | 1.160 |
| Evenness ($e^H/S$) | 0.967 | 0.819 | 0.791 | 1.005 | 0.772 | 0.771 | 1 | 0.488 | 0.532 |

**Figure 2.**
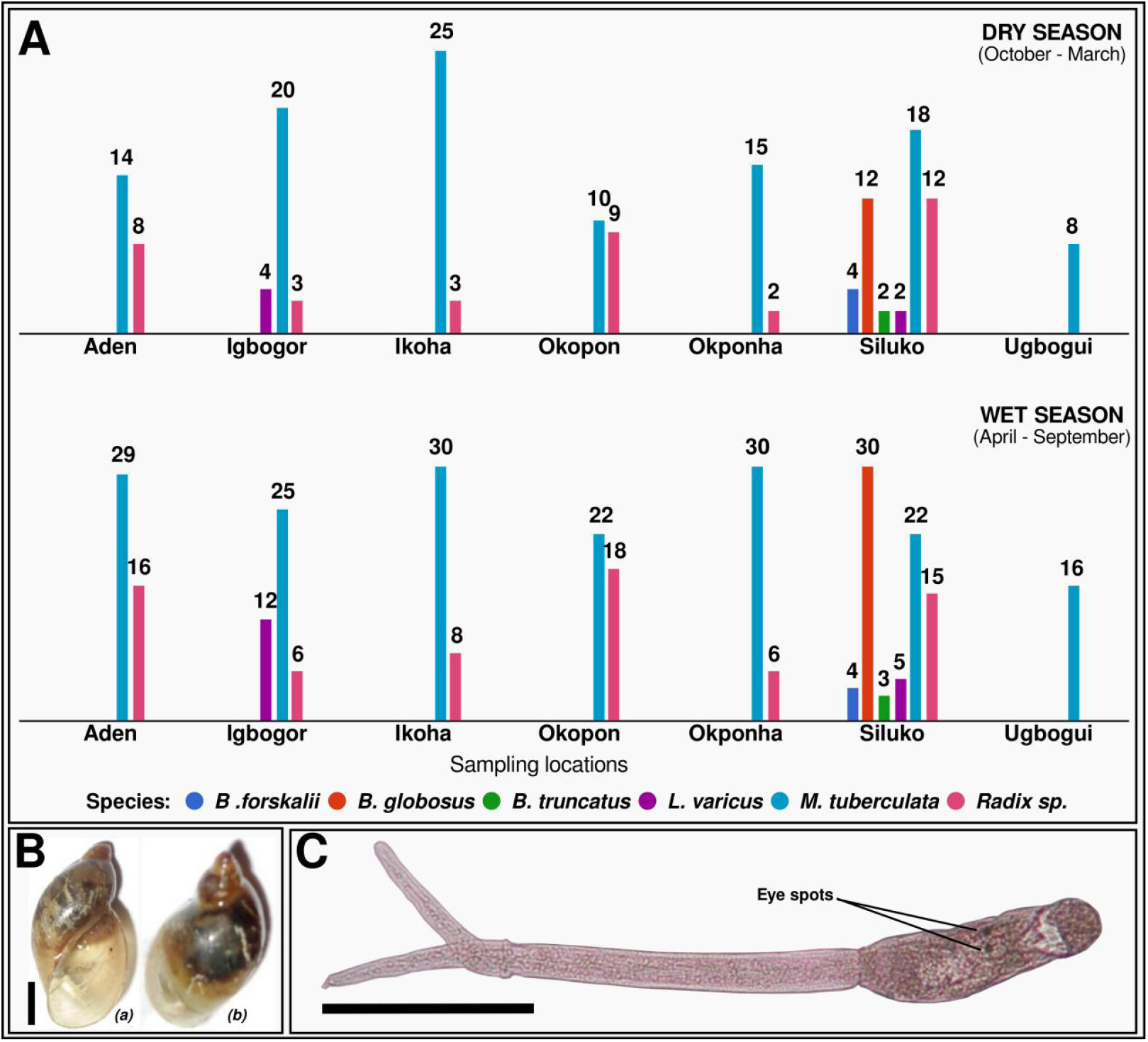
A, Bar chart of the seasonal distributions of freshwater snails collected in Ovia South-West, Edo state, Nigeria. The number of individual snail of each species are shown on top of each bar. B, *Radix* sp. collected in Siluko infected with brevifurcate-apharyngeate distome furcocercaria. Scale bar: 2 mm (a) Apetural, and (b) Lateral views. C, Brevifurcate-apharyngeate distome furcocercaria from *Radix* sp. collected in Siluko. Scale bar: 200 μm. Note the prominent eye spots.

### Snail Diversity Indices

The Shannon diversity index (H) of the snails from the various sampled locations in Ovia South-West LGA of Edo State ranged from 0 to 1.532 (Table 2). Siluko recorded the highest species diversity (H = 1.532 and D = 0.248), followed by Igbogor (H = 0.899, D - 0.475). The lowest species diversity was at Ugbogui, where only *M. tuberculata* was recorded (H = 0 and D = 1). Kruskal-Wallis test indicated no statistical significance in snail diversity between the sites at α = 0.05 (test statistics H = 6, degree of freedom = 6; critical value = 12.59). The evenness of snails across the sampling locations was moderately to perfectly even, with a range between 0.771 and 1.005. Snail collection during the wet season had a slightly higher diversity and evenness compared to the dry season (Table 2). However, the Mann-Whitney U test at a significance level of 0.05 indicated no significant difference in diversity between the dry and wet seasons (test statistics U = 14.5, critical value = 5).

### Trematode Infection in Snails

The known vector snails of schistosomiasis (*B. forskalii, B. globosus*, and *B. truncatus*) collected during this study were uninfected with *Schistosoma* cercariae. Apart from a single *Radix* sp. (Figure 2B) collected in Siluko, all other snail species were uninfected. The infected *Radix* from Siluko shed brevifurcate-apharyngeate distome furcocercaria trematode with distinct eye spots (Figure 2C).

## Discussion

Ovia South-West LGA of Edo State is a known endemic location for urogenital schistosomiasis in Nigeria (Adeyemi *et al*., 2014; Noriode *et al*., 2017; Enabulele *et al*., 2021). Before this study, only Igbinosa *et al*. (2015) had investigated the mollusc diversity in the LGA and reported three species of freshwater snails (*Gabbiella humerosa, L. varicus*, and *M. tuberculata*). More diversity of freshwater snails (six species) is reported in the present study when compared with Igbinosa *et al*. (2015). Four genera of snails - *Bulinus, Lanistes, Melanoide*s, and *Radi*x, were recorded during a year-long sampling of rivers in seven localities in Ovia southwest. Based on the morphological appearance of shells, five of the snails (*B. forskalii, B. globosus, B. truncatus, L. varicus*, and *M. tuberculata*) were identified at the species level (Table 1). One snail could only be tentatively identified to the genus level as *Radix* sp. (Figure 2B).

The diversity of freshwater snails reported is consistent with the species often reported in other molluscan surveys in Nigeria (Ndifon and Ukoli, 1989; Salawu and Odaibo, 2014; Anorue *et al*., 2024). *Melanoides tuberculata* was the most abundant species across all locations (n = 284, 60.68%). It was the only species occurring in all seven sampled locations and dominated (Table 1). A similar abundance of *M. tuberculata* has been reported in other locations in Nigeria (Ndifon and Ukoli, 1989; Oloyede *et al*., 2016; Oladejo *et al*., 2021; Oso and Odaibo, 2021). The combination of several biological traits, ecological adaptability, and human-mediated factors have contributed to *M. tuberculata* being a dominant species in many freshwater ecosystems where they are native or invasive (Pointier *et al*., 1992).

The occurrence of *B. forskalii, B. globosus*, and *B. truncatus* confirms that these known snail hosts of *S. haematobium* are possibly responsible for transmitting urogenital schistosomiasis in Ovia South-West LGA of Edo State. Snail hosts for *S. mansoni, Biomphalaria*, were not recorded and may explain why intestinal schistosomiasis has remained unreported in the LGA. Among the *Bulinus* species (*B. forskalii, B. globosus, B. truncatus*) recorded, *B. globosus* (n = 42) was the most abundant species (Table 1). This species is widely recognized as the intermediate host for *S. haematobium* in Nigeria and is often reported as the dominating *Bulinus* species in water bodies in communities endemic for urogenital schistosomiasis (Odaibo *et al*., 2004; Odeniran *et al*., 2020; Oso and Odaibo, 2021; Anorue *et al*., 2024). Only in Siluko were all the *Bulinus* species recorded, both dry and wet seasons (Figure 2A). It is unclear why Siluko was the only site for the *Bulinus* species. However, the physicochemical properties of water bodies, either influenced by anthropogenic activities or natural habitat, and competitor snails such as *M. tuberculata* are known drivers of *Bulinus* occurrence in a freshwater ecosystem (Pointier and Jourdane, 2000; Oso and Odaibo, 2021; Kagabo *et al*., 2024; Tabo *et al*., 2024).

Three species of *Lanistes, L. libycu*s, *L. ovum*, and *L. varicus*, have been reported in Nigeria (Ndifon and Ukoli, 1989; Salawu and Odiabo, 2014; Olorunniyi and Olofintoye, 2024). *Lanistes varicus* was recorded in two locations (Igbogor, n = 16 and Siluko, n = 7) (Table 1). In contrast to our findings, Igbinosa *et al*. (2015) reported *L. varicus* as the most abundant species (44.6%, n = 258) across four sample locations different from the sites we sampled in Ovia southwest in Edo State. Experimental studies using Ghanian strains of *L. varicus* indicated their ability to serve as potential biological control for *B. truncatus* and *Biomphalaria pfeifferi* (Anto *et al*., 2005; Anto and Bimi, 2017). It is uncertain if such experimental findings apply in field conditions and impact the occurrence of *Bulinus* in Siluko and Igbogor. Although Igbinosa *et al*. (2015) didn’t record any *Bulinus* species in all the locations where *L. varicus* occurred, the presence of the species seems not to have impacted the occurrence of *B. globosus* (50.5%) and *B. truncatus* (3.7%) in Ebonyi, southeastern Nigeria (Anorue *et al*., 2024).

The unidentified species of *Radix* (Figure 2B) recorded was present in six of the seven sampling locations (Table 1). Based on the shell morphology, the *Radix* species was identified as *Lymnaea natalensis* (personal communication in 2014 with Professor Odaibo AB of the University of Ibadan, Nigeria). *Lymnaea natalensis* Krauss 1848 was reclassified to *Radix natalensis* as part of broader taxonomic revisions in the late 20th and early 21st century when molecular phylogenetic studies revealed that the genus *Lymnaea* was not monophyletic (Glöer, 2002; Correa *et al*., 2010; Vinarski, 2013). The species has been widely reported in Nigeria (Ndifon and Ukoli, 1989; Ofoezie, 1999; Abe *et al*., 2016; Oladejo *et al*., 2021) and is primarily recognized as an intermediate host for *Fasciola gigantica* in Africa (Brown, 1994; Vinarski *et al*., 2019; Malatji *et al*., 2019).

We hesitate to identify the *Radix* from Ovia southwest as *R. natalensis* because of the similar shell morphology with *Pseudosuccinea columella*, an invasive snail that has been reported in several countries in Africa (Brown, 1994; Malatji *et al*., 2019; Jones *et al*., 2024). At the time of our study, we lacked resources for thorough conchological or anatomical analysis including the ability to take high-resolution photographs of the shells to identify the characteristic surface spiral rows of short transverse grooves which differ from the spiral ridges characteristic of *P. columella* (Brown, 1994; Jones *et al*., 2024). We also lacked molecular markers to characterize the specimen for accurate identification (Correa *et al*., 2010).

Out of the four hundred and sixty-eight snails screened for trematode, only a single *Radix* sp. collected in Siluko was infected. While it is not uncommon for there to be the absence or low prevalence of trematode infection in snails in a freshwater habitat (Agi, 1995; Mafiana and Beyioku, 1998; Adediran *et al*., 2014), several factors could be responsible. Low snail density, low presence or absence of appropriate infected intermediate, definitive or reservoir hosts, and the physico-chemical properties of the water body are factors that can determine the level of infection of snail populations with trematodes (Buck *et al*., 2017; Hobart *et al*., 2022; Song and Proctor, 2020). It is also possible we missed infected snails at the prepatent stage, and screening techniques such as snail crushing or molecular approaches like xenomonitoring could have been useful (Schols *et al*., 2020; Pennance *et al*., 2020; Gaye *et al*., 2024).

The trematode recovered from a single *Radix* sp. in Siluko (Figure 2C) was a brevifurcate-apharyngeate distome furcocercarial type (Frandsen and Christensen, 1984). Although Siluko is endemic for urogenital schistosomiasis, we doubt that the cercaria is a *Schistosoma* sp. because of the conspicuous eye spots. In Africa, brevifurcate-apharyngeate distome furcocercaria with eye spots are typically either species in the family *Spirorchiidae* (blood flukes of reptiles) or blood flukes of birds such as *Trichobilharzia* and *Gigantobilharzia* (Frandsen and Christensen, 1984). Although morphological characteristics of the cercaria from the *Radix* sp. suggest it is not a *Schistosoma* species, molecular phylogenetic analysis will provide definitive identification (Blasco-Costa *et al*., 2016).

## Conclusion

A year-long malacological survey of freshwater bodies in Ovia South-West Local Government Area in Edo State, Nigeria, revealed the presence of six snail taxa. Five species, *B. forskalii, B. globosus, B. truncatus, L. varicus*, and *M. tuberculata*, were identified, and one only to the genus *Radix* based on shell morphology. Accurate identification of the *Radix* species will require detailed conchological, anatomical, and molecular analysis because of its similarity to *R. natalensis* and *P. columella*. Although Ovia South-West is endemic for urogenital schistosomiasis, none of the collected *Bulinus* screened for schistosome cercariae were infected. The *Radix* sp. collected in Siluko shed brevifurcate-apharyngeate distome furcocercariae. The conspicuous eye spots on the cercaria suggest it is not a *Schistosoma* sp. commonly responsible for schistosomiasis. Definitive identification of the brevifurcate-apharyngeate furcocercaria infecting the *Radix* sp. in Ovia South-West will require molecular phylogenetics analysis.

## Acknowledgements

The authors acknowledged the assistance of Prof. A. B Odaibo of the Department of Zoology, University of Ibadan, for snail identification.

## Conflicts of interest

The author declare that they have no conflict of interest

## Notes

### Competing Interest Statement

The authors have declared no competing interest.

